# Targeting Acid Ceramidase Ameliorates Fibrosis in Mouse Models of Nonalcoholic Steatohepatitis

**DOI:** 10.1101/2022.07.10.499388

**Authors:** Amy Yu, Carson Cable, Mahbubul H. Shihan, Sachin Sharma, Aras N. Mattis, Izolda Mileva, Yusuf A. Hannun, Caroline C. Duwaerts, Jennifer Y. Chen

## Abstract

Nonalcoholic fatty liver disease (NAFLD) is a common cause of liver disease worldwide, and is characterized by the accumulation of fat in the liver. Nonalcoholic steatohepatitis (NASH), an advanced form of NAFLD, is a leading cause of liver transplantation. Fibrosis is the histologic feature most associated with liver-related morbidity and mortality in patients with NASH, and treatment options remain limited. In previous studies, we discovered that acid ceramidase (aCDase) is a potent antifibrotic target using human hepatic stellate cells (HSCs) and models of hepatic fibrogenesis. Using two dietary mouse models, we demonstrate that depletion of aCDase in HSC reduces fibrosis without worsening metabolic features of NASH, including steatosis, inflammation, and insulin resistance. Consistently, pharmacologic inhibition of aCDase ameliorates fibrosis but does not alter metabolic parameters. The findings suggest that targeting aCDase is a viable therapeutic option to reduce fibrosis in patients with NASH.

## 2) INTRODUCTION

Characterized by progressive matrix stiffening, tissue fibrosis is a leading cause of morbidity and mortality^1^. Despite steady progress in basic, translational, and clinical research of fibrogenesis, there remain limited treatment options for patients. Organ transplantation is an effective option for end-stage disease, but is limited by donor organ availability. This underscores the need for new therapies for patients with fibrotic diseases.

The burden of end-stage liver disease has risen significantly due to the increasing global prevalence of nonalcoholic fatty liver disease (NAFLD). NAFLD is characterized by the accumulation of lipids within hepatocytes and is associated with features of metabolic syndrome such as obesity, insulin resistance, and hyperlipidemia. NAFLD represents a spectrum of liver disease that can lead to progressive nonalcoholic steatohepatitis (NASH), fibrosis, and ultimately hepatocellular carcinoma and liver failure. Among patients with NASH, fibrosis is the histologic measure that predicts liver-related mortality^2^, further highlighting the need for antifibrotic therapies.

Hepatic stellate cells (HSCs) drive liver fibrosis, and our prior studies identified a new antifibrotic target, acid ceramidase (aCDase)^3^, an enzyme responsible for ceramide hydrolysis. We demonstrated that targeting aCDase promotes HSC inactivation^3^. In additional studies, we showed that genetic deletion in HSCs or pharmacologic inhibition of aCDase ameliorates fibrosis in mouse models. Mechanistically, we illustrated that targeting aCDase inhibits YAP/TAZ activity by potentiating its proteasomal degradation. We also demonstrated that aCDase inhibition reduces fibrogenesis in human fibrotic precision-cut liver slices. Consistently, patients with advanced fibrosis have increased aCDase expression compared to those with mild fibrosis^4^. Furthermore, a signature of the genes most downregulated by ceramide, the ceramide responsiveness score (CRS), identifies patients with advanced fibrosis who could benefit from aCDase targeting^4^.

Given the rising prevalence of NAFLD and the lack of available therapies, we aimed to determine how targeting aCDase regulates fibrosis and metabolic parameters in mouse models of NASH. These studies are particularly relevant, as others have shown that ceramide species can contribute to insulin resistance and steatosis^5-10^, which would complicate NASH treatment. We previously observed that genetic depletion of aCDase in HSCs ameliorates fibrosis development without altering steatosis in one model of NASH, the choline-deficient L-amino acid-defined, high-fat diet (CDAHFD) model^4^. Here, we aimed to characterize the effect of targeting aCDase on metabolic parameters in the CDAHFD model and in a second dietary model of NASH, the Fructose, Palmitate, Cholesterol (FPC) model. In this study, we demonstrate that genetic deletion in HSCs or pharmacologic inhibition of aCDase reduces fibrosis but does not worsen metabolic parameters of NASH. This work highlights the therapeutic potential of aCDase targeting in patients with NASH.

## 3) MATERIALS AND METHODS

Animal experiments were approved by the Institutional Animal Care and Use Committee at the University of California, San Francisco. All animals received humane care according to the criteria outlined in the *Guide for the Care and Use of Laboratory Animals of the National Academy of Sciences*.

To generate the HSC depletion of aCDase, we crossed *Asah1*^flox/flox^ (provided by Lina Obeid)^11^ with *Pdgfrb*-Cre^12^ (*Asah1*^cko^, cACKO). Control mice were *Asah1*^flox/flox^ lacking *Pdgfrb*-Cre. In the choline-deficient L-amino acid-defined, high-fat diet (CDAHFD) model, 6-to 8-week-old male mice received either normal chow (PicoLab Rodent Diet 20; LabDiet #5053) or CDAHFD (L-amino acid diet with 60 kcal% fat with 0.1% methionine without added choline; Research Diets A06071302) *ad libitum* for 14 weeks. In the Fructose, Palmitate, and Cholesterol (FPC) model, 6-to 8-week-old sex-matched mice received either normal chow or FPC (Fructose, Palmitate, and Cholesterol; Envigo TD 160785) supplemented with high fructose drinking water (45:55 fructose:glucose) *ad libitum* for 16 weeks^13^. Mice were weighed and food consumption measured once a week. Weekly food intake was measured by monitoring the weight difference between added food and the remaining food.

For the therapeutic B13 experiment, male C57BL/6J mice (age 6-8 weeks, Jackson Laboratory, Bar Harbor, ME) received FPC diet *ad libitum* for 9 weeks. Mice then received either 50 mg/kg B13 or vehicle 5 days/week by IP for 3 weeks (total weeks on the FPC diet = 12 weeks).

A terminal blood collection was performed by cardiac puncture. Livers were weighed and were subsequently fixed in formalin, 4% paraformaldehyde, or snap frozen for further analysis. Mice were fasted for 4 hours prior to sacrifice.

### Serum Analysis

Harvested blood was processed for collection of serum. Serum was used to measure ALT and total cholesterol at Zuckerberg San Francisco General Hospital. The insulin ELISA was performed on mouse serum according to the manufacturer instructions for the “Low Range Assay” of the kit (Crystal Chem, Ultra Sensitive Mouse Insulin ELISA Kit #90080).

### Hepatic Triglycerides

Lipids were extracted from liver tissue using the Folch method^14^. Liver triglycerides were quantified as described previously^15^.

### Sphingolipid Analysis

Sphingolipid analysis was performed as described previously using LC/MS/MS^3^.

### Glucose Tolerance Test

Mice were fasted for 5 hours before blood was collected from the base of the tail. Glucose was measured using a glucometer (Accu-Chek Performa Nano #06454283056). Blood glucose was measured at start of test (0min time-point). Mice then received 20% glucose solution via intraperitoneal (IP) injection (10uL per gram of weight), and glucose measurements were taken at the following time points after IP injection: 15, 30, 60, 90, and 120 minutes.

### Histology, Sirius Red Staining

Formalin-fixed samples were embedded in paraffin, cut in 5 μm sections, and stained with hematoxylin & eosin (H&E) by Peninsula Histopathology Laboratory. Sirius red staining was performed as previously described^12^. The collagen proportional area (CPA) was morphometrically quantified on Sirius red-stained sections with ImageJ as previously described^12^. Slides were evaluated by a blinded expert UCSF liver pathologist (A.N.M) for steatosis and lobular inflammation using a histological scoring system for nonalcoholic fatty liver disease^16^.

### Statistics

Statistical analysis was performed using GraphPad Prism 8 with unpaired two-sided student t-tests, one-way ANOVA with Tukey’s method for multiple comparisons, or Kruskal-Wallis test with Dunn’s multiple comparisons test. Statistical significance was defined as P < 0.05.

## 4) RESULTS

### HSC depletion of aCDase does not worsen metabolic features in the CDAHFD model of NASH

In previous studies, we demonstrated that HSC deletion of aCDase reduces fibrosis development in the choline-deficient L-amino acid-defined, high-fat diet (CDAHFD) model of NASH^4^. Among mice fed CDAHFD, we did not observe significant differences in the development of steatosis, lobular inflammation, or hepatic triglyceride levels between mice with HSC deletion of aCDase and their Cre-negative littermate controls^4^.

Here, we explored how HSC deletion of aCDase regulates metabolic parameters of NASH in the CDAHFD model (Fig 1A). There was, as expected, an overall significant decrease in weight gain for mice on CDAHFD combined with a significant decrease in food intake (Fig 1B, C). Also as expected with this dietary model, the mice experienced no insulin resistance as demonstrated by a normal glucose tolerance test (Fig 1D). At the time of sacrifice, mice receiving CDAHFD experienced significantly decreased fasting blood glucose and insulin levels compared to those receiving standard chow diet, but there were no significant differences between mice receiving CDAHFD (Fig 1E, F). This suggests that although the CDAHFD model produces fibrosis, steatosis, lobular inflammation, and increases in hepatic triglycerides, this model does not recapitulate the weight gain or insulin resistance observed in patients with NASH and thus represents a suboptimal mouse model.

**Figure 1.**
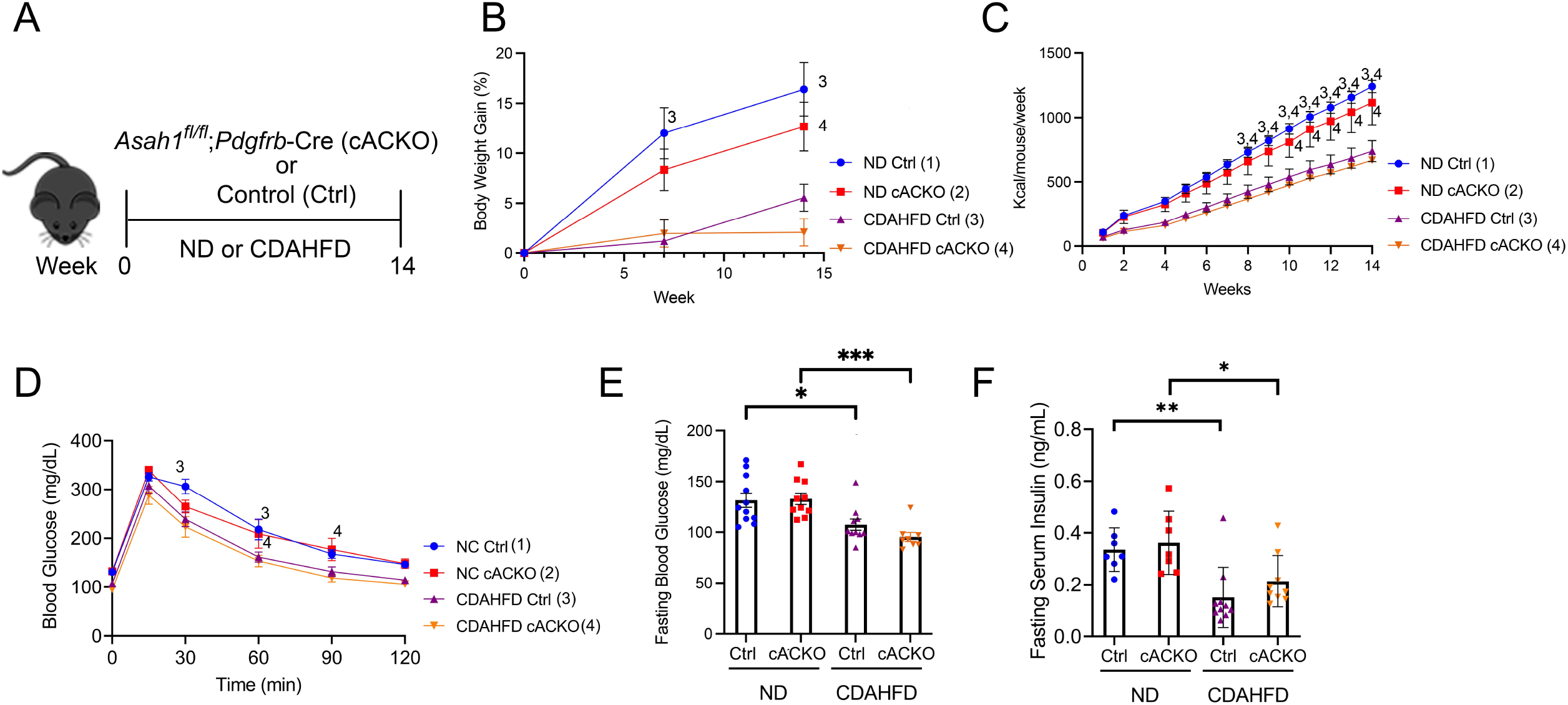
Hepatic stellate cell-specific acid ceramidase knockout mice on a CDAHFD demonstrate no metabolic differences compared with their wild-type counterparts. (A) Experimental schematic. Male and female hepatic stellate cell-specific acid ceramidase knockout mice (cACKO) or control (Ctrl) mice were fed either normal diet (ND) or CDAHFD for 14 weeks. (B) Body weight percent over time. (C) Food intake in Kcal/mouse/week. (D) Glucose tolerance test. (E) Fasting blood glucose levels measured at 14 wk. (F) Fasting serum insulin levels measured at 14 wk. n = 8-11 mice per group. *p<0.05, **p<0.01, ^1-4^p<0.05 to the corresponding group number.

### HSC depletion of aCDase does not worsen metabolic features in the FPC model of NASH

To address the limitations of the CDAHFD model, we next utilized the FPC model of NASH, which has been shown to induce insulin resistance after 16 weeks.^13^ The FPC model was previously studied in male mice^13^, and we sought to decipher whether there were sex-specific differences in the development of fibrosis and features of metabolic syndrome.

Female mice receiving the FPC diet experienced significant increases in hepatic steatosis and weight gain compared to mice receiving the standard chow diet (Fig 2J, L), and a similar trend was observed among male mice (Fig. 2D, F). Among male and female mice receiving the FPC diet, there were no significant differences between the conditional knockout mice and control mice with respect to ALT, steatosis, inflammation, and weight gain (Fig 2B-F, 2H-L).

**Figure 2.**
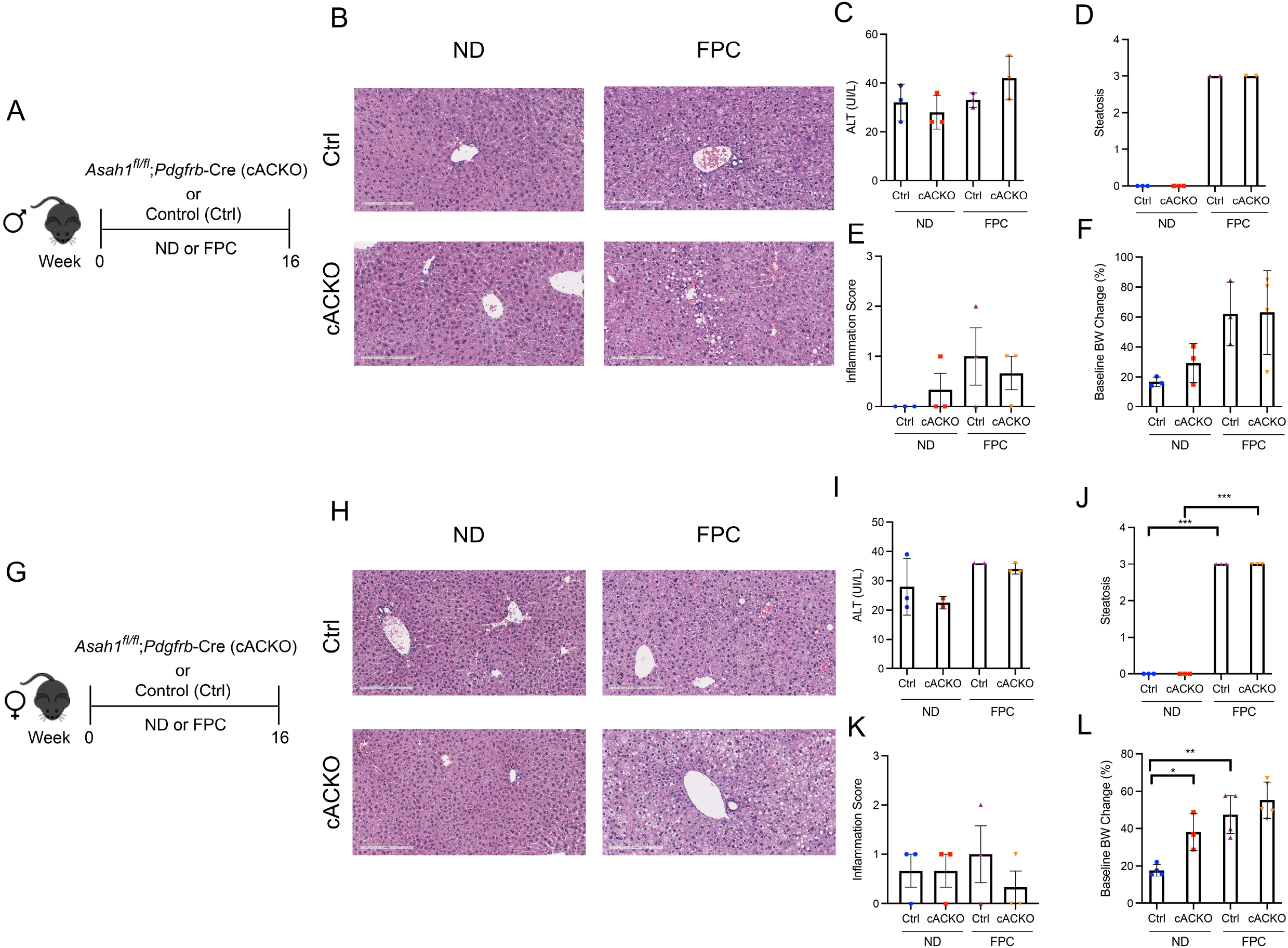
Hepatic stellate cell-specific acid ceramidase knockout mice on a FPC diet demonstrate no liver or metabolic differences compared with their wild-type counterparts. (A, G) Experimental schematic. Male and female hepatic stellate cell-specific acid ceramidase knockout mice (cACKO) or control (Ctrl) mice were fed either normal diet (ND) or Fructose Palmitate Cholesterol diet (FPC) for 16 weeks. (B, H) Representative photomicrograph of H&E-stained liver sections. (C, I) Measured serum ALT levels. (D, J) Liver steatosis graded by a blinded pathologist. (E, K) Inflammation score graded by a blinded pathologist. (F, L) Body weight (BW) percent change at 16wk post-diet from start weight. n = 2-6 mice per group. *p<0.05, **p<0.01, ***p<0.001.

### Male mice with HSC depletion of aCDase have a trend towards decreased fibrosis compared to control male mice

Consistent with our prior data using the CDAHFD model^4^, male mice with HSC deletion of aCDase had significantly decreased fibrosis compared to control mice receiving FPC as measured by collagen proportional area (Fig 3A-C). We did not observe significant differences among female mice receiving the FPC diet (Fig 3D-F), suggesting there may be sex differences that modulate this response.

**Figure 3.**
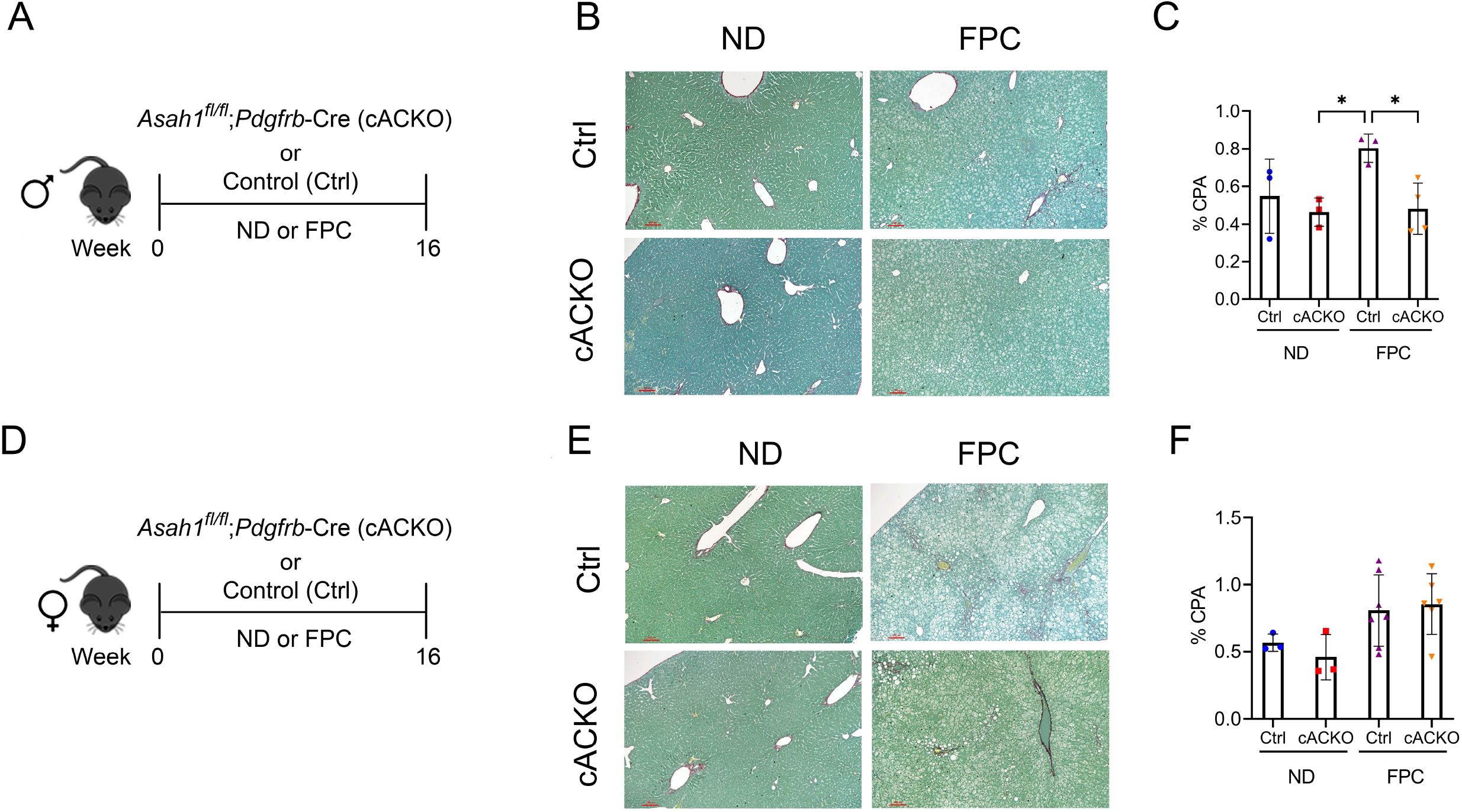
Male hepatic stellate cell-specific acid ceramidase knockout mice on a FPC diet develop significantly less fibrosis than wild-type counterparts. (A,D) Experimental schematic. Male or female mice of either hepatic stellate cell-specific acid ceramidase knockout (cACKO) or their respective Cre-negative littermate controls (Ctrl) received either normal diet (ND) or Fructose Palmitate Cholesterol (FPC) for 16 wk. (B,E) Representative photomicrograph of Sirius red staining for either male (top) or female (bottom) mice. (C,F) Quantification of positive collagen proportional area (Sirius red staining, % CPA) for males (top) or females (bottom). n = 3-6 mice per group. *p<0.05

### HSC depletion of aCDase does not worsen metabolic parameters of NASH in male mice

Intrigued by these sex-specific differences, we next aimed to characterize metabolic parameters in the conditional knockout mice according to sex. Among male mice, the FPC diet induced significant increases in hepatic triglycerides, liver proportional weight, and serum cholesterol, but there were no significant differences between FPC-fed conditional knockout mice and control mice (Fig 4A-D). We observed consistent findings among the female mice (Fig S1A-C).

**Figure 4.**
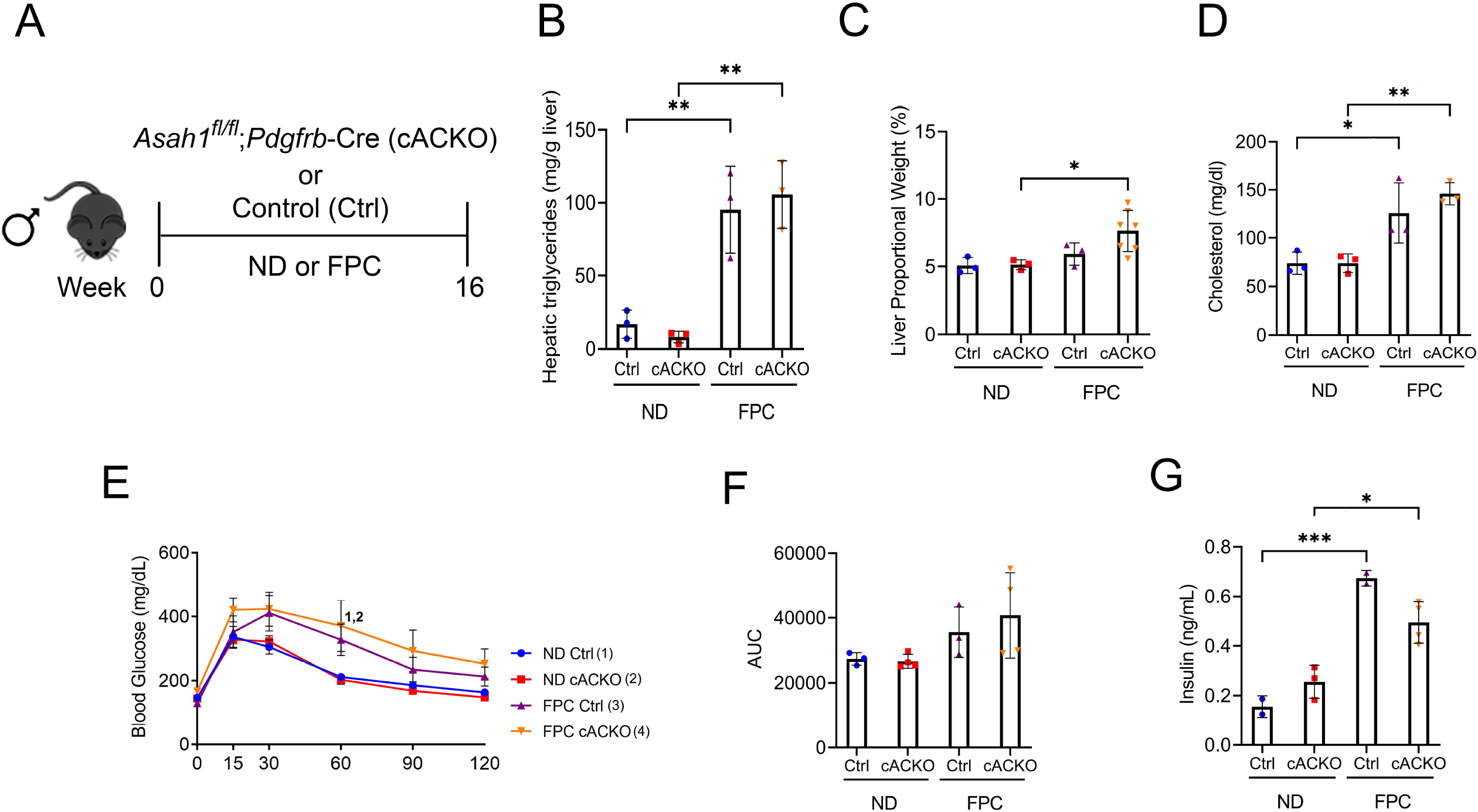
Male hepatic stellate cell-specific acid ceramidase knockout mice demonstrate no metabolic differences from their wild-type counterparts. (A) Experimental schematic. Male hepatic stellate cell-specific knockout (cACKO) or control (Ctrl) mice were fed either a normal diet (ND) or Fructose Palmitate Cholesterol (FPC) diet for 16 wk. (B) Measured hepatic triglycerides. (C) Liver to body proportional weight in percent. (D) Measured serum cholesterol levels. (E-F) Glucose tolerance test was performed at 14 wk and area under the curve (AUC) was determined. (G) Measured serum insulin levels. n = 3-4 mice per group. *p<0.05, **p<0.01, ***p<0.001, ^1-4^p<0.05 to the corresponding group number.

In contrast to the CDAHFD model, we observed increases in glucose intolerance among male mice receiving FPC compared to standard chow diet as measured by a glucose tolerance test (GTT). There were no significant differences among FPC-fed mice (Fig 4E, F). Interestingly, we observed a trend towards a decrease in fasting serum insulin among FPC-fed conditional knockout male mice compared to control mice (Fig 4G), which was not seen in female mice (Fig S1F).

To measure changes in ceramide subspecies, we performed sphingolipid analysis of liver tissues (Fig S2). This analysis was performed on male mice as we observed a decrease in fibrosis in this group only. We observed that the FPC diet significantly increased C20:1 and C22:1 compared to normal diet. We also observed that the FPC significantly decreased C18. However, there were no significant differences in ceramide subspecies between mice receiving normal or FPC diet (Fig S2).

### Pharmacologic inhibition of aCDase ameliorates fibrosis and does not worsen metabolic parameters

Our studies thus far have analyzed the role of HSC deletion of aCDase in mouse models of NASH, but the impact of systemic aCDase inhibition has not been elucidated. To further investigate the effect of targeting aCDase for the treatment of NASH, we utilized a pharmacologic inhibitor, B13. B13 has been validated as an aCDase inhibitor in other cell and model systems^17-22^, including by our group^4^. We next explored how B13 treatment regulates fibrogenesis in the FPC model. We established the NASH phenotype by providing 6-8 week old male C57BL/6 mice FPC diet for 9 weeks. Mice then received either B13 or vehicle for an extra 3 weeks while still on diet (total weeks on FPC diet = 12 weeks) (Fig 5A). Given our observation of sex-specific differences in the development of fibrosis using the FPC model, we included only male mice in this analysis.

**Figure 5.**
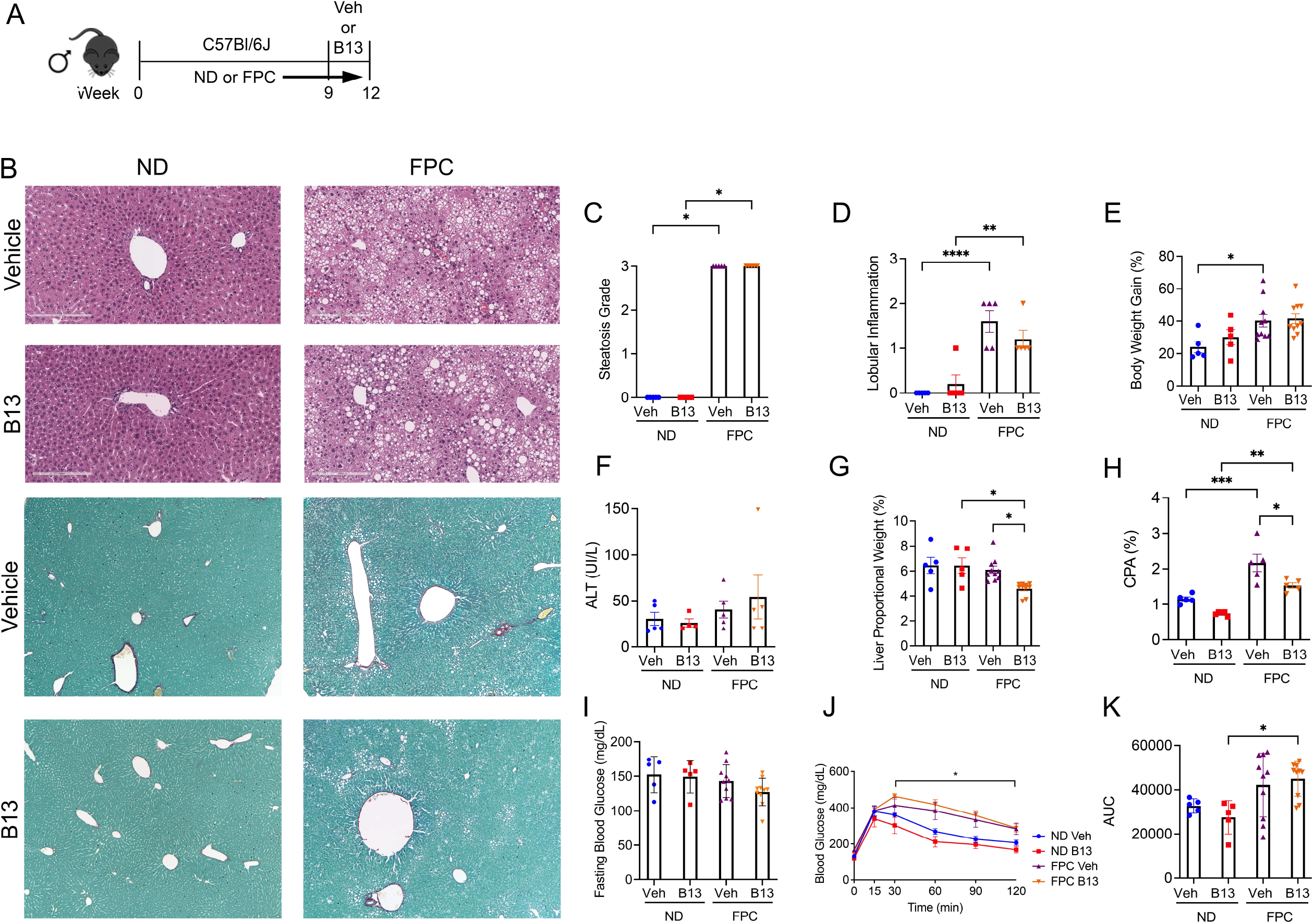
Male wild-type mice on FPC diet and administered an acid ceramidase inhibitor developed less fibrosis than vehicle control mice. (A) Experimental schematic. Male wildtype mice received either normal diet (ND) or Fructose Palmitate Cholesterol (FPC) diet for a total of 12 wks. During the last 3 weeks of feeding, they also received either a small molecule acid ceramidase inhibitor (B13, dosed at 50 mg/kg by intraperitoneal injection five times per week) or vehicle control (Veh). (B) Representative photomicrograph of H&E and Sirius-red stained liver sections. A pathologist graded (C) liver steatosis and (D) lobular inflammation. (E) Percent body weight gain from start. (F) Measured serum ALT levels. (G) Liver to body weight proportional weight in percent. (H) Sirius red staining was quantified (collagen proportional area, CPA). (I) Fasting blood glucose levels were measured at week 12. (J-K) Glucose tolerance test was performed at week 10 and area under the curve (AUC) was calculated. n = 5-10 mice per group. *p<0.05, **p<0.01, ***p<0.001, ****p<0.0001.

Receipt of FPC diet significantly increased steatosis, lobular inflammation, and weight gain, but there were no significant differences between FPC-fed mice receiving B13 or vehicle (Fig 5B-E). We did not observe significant changes in ALT level (Fig 5F). The liver proportional weight was lower among FPC-fed mice receiving B13 compared with vehicle (Fig 5G).

Notably, in the FPC model, B13 significantly reduced fibrosis as measured by collagen proportional area (Fig 5B, H). The FPC diet induced a significant increase in glucose intolerance among B13-treated mice (Fig 5J, K). However, among FPC-fed mice receiving B13 or vehicle, there were no significant differences in glucose intolerance (Fig 5J, K).

Furthermore, we performed sphingolipid analysis of liver tissues (Fig S3). We observed that the FPC diet significantly increased several subspecies including C16, C20:1, C22:1, C26, dhC18, dhC18:1, dhC20, dhC20:1, dhC22, and dhC22:1. Interestingly, several ceramide subspecies significantly decreased with the FPC diet, such as C18 and C24:1. Among mice fed normal diet, B13 treatment significant reduced C24:1 and C26:1. Among mice fed FPC diet, there was a significant decrease in C20:1 ceramide with B13 treatment compared to vehicle. Taken together, our findings demonstrate that pharmacologic inhibition of aCDase reduces fibrosis without altering metabolic parameters in the FPC model of NASH.

## 5) DISCUSSION

The burden of nonalcoholic fatty liver disease (NAFLD) is rising globally, and there are limited treatment options available for patients. In particular, there are no treatment options targeting fibrosis, which drives liver-related morbidity and mortality^2^. Our studies suggest that genetic deletion of aCDase in HSCs or pharmacologic inhibition of aCDase ameliorates fibrosis without worsening metabolic features of NASH. This suggests that targeting aCDase represents an effective strategy to reduce fibrosis in this disease setting.

Ceramides constitute a family of sphingolipids that consist of sphingosine linked to a fatty acid, and are generated through several pathways, including de novo synthesis from serine and palmitate, sphingomyelin hydrolysis, or recycling of sphingosine. Ceramide species can regulate diverse cellular behavior^23^, and ceramide metabolism has been implicated in the pathogenesis of insulin resistance and steatosis^5-10^, though the literature is conflicting. Inhibition of de novo synthesis, including targeting dihydroceramide desaturase (DES1), mitigates NASH phenotypes^6,24,25^. Human lipidomic studies have also demonstrated a correlation between insulin resistance and hepatic ceramides^26,27^. However, targeting other aspects of ceramide metabolism has had a varying impact. Overexpression of acid ceramidase in hepatocytes or adipose tissue improves insulin sensitivity in mice receiving a high-fat diet^10^, but deficiency in alkaline ceramidase 3 alleviates inflammation and fibrosis in a mouse model of NASH^28^. Furthermore, ceramide subspecies have different cell-specific functions: for example, C_16_-ceramide activates apoptosis, whereas C_22_-ceramide and C_18:1_-ceramide inhibit apoptosis in hepatocytes^29,30^. A recent study demonstrated that short-chain C_6_-ceramide promotes anti-oxidant signaling in a mouse model of NASH^31^. Thus, it is overly simplistic to stipulate that all ceramides are pathologic in NASH or other disease settings.

Our prior studies demonstrate a pivotal role of aCDase in regulating HSC activity and hepatic fibrosis in several systems, including models of NASH^3,4^. For example, treatment with B13 reduced fibrogenesis in precision-cut fibrotic liver slices from CDAHFD-fed rats, and mice with deletion of aCDase in HSCs experienced decreased fibrosis development in this dietary model^4^. Furthermore, a signature consisting of the top genes downregulated by ceramide, the Ceramide Responsiveness Score, was significantly increased in NAFLD patients with advanced compared to mild fibrosis. This suggests that the transcriptional response associated with increasing fibrosis could be reversed with aCDase inhibition in patients with NAFLD^4^. Here, our studies demonstrate that the metabolic parameters of NASH including glucose tolerance, hepatic triglycerides, and steatosis are not significantly altered with HSC depletion or pharmacologic inhibition of aCDase. Taken together, our findings highlight that targeting aCDase has significant antifibrotic effects that are dissociated from metabolic parameters. This suggests that aCDase acts downstream of the initial metabolic perturbations that lead to NASH and most likely at the level of HSCs, which transmit these proximal changes into a fibrotic response.

This work has several limitations. We intentionally selected a therapeutic intervention in which the NASH phenotype was established before treatment with the aCDase inhibitor B13, as this mimics treatment of patients with established disease. However, this timing does not allow for analysis of how targeting aCDase modulates development of steatosis and/or steatohepatitis. We also did not explore how aCDase targeting regulates adipose tissue and insulin signaling. We performed sphingolipid analysis on mouse hepatic tissues in 2 experiments using the FPC diet. We observed that the FPC diet induces changes in different ceramide subspecies, including decreases in several subspecies, further highlighting the diversity of ceramides. Interestingly, B13 treatment modulated levels of C24:1 and C26:1 in mice fed a normal diet and C20:1 in mice fed FPC. We also want to highlight that not all ceramides were measured in the analysis. Of note, we previously measured sphingolipids in mice receiving systemic B13 in a carbon tetrachloride model of fibrosis: in this study, B13 increased two specific subspecies, C_26_-ceramide and C_26:1_-ceramide^4^, which was not observed in these studies with the FPC model. Additional studies are underway to measure cell-specific changes in sphingolipids, including short-chain ceramides, and to determine whether compensation by other pathways involved in ceramide metabolism occurs with aCDase deletion. Furthermore, we selected two dietary models of NASH, but other dietary or genetic models will be important to explore in future studies.

Our findings also highlight potential sex differences in response to the FPC model. Human NASH studies as well as studies on hepatocellular carcinoma (HCC) incidence have demonstrated a significant difference in incidence between males and females. Studies have illustrated a clear link between estrogen and the decreased occurrence of HCC^32,33^. The same phenomenon holds true for NASH. Studies have shown that NASH is more prevalent in males and post-menopausal women than pre-menopausal women^34,35^. However, studies investigating NASH fibrosis have been inconclusive in a potential role for sex in its incidence^**3**6-38^. In our studies using the FPC model, we observed significant increases in serum insulin levels among male mice receiving the FPC diet compared to the standard chow diet. This difference was not observed among female mice, which is consistent with prior studies demonstrating that female mice are less prone to develop insulin resistance^39^. We acknowledge that the studies were underpowered to comprehensively analyze differences by sex and future studies are needed to characterize differences with the FPC model.

In summary, our studies suggest that targeting aCDase in two dietary models of NASH reduces fibrogenesis without worsening metabolic features. Our findings suggest that this strategy of targeting aCDase may add to the armamentarium of antifibrotic therapies for patients with NASH as well as other types of chronic liver disease.

## Figure Legends

**Supplemental Figure 1. Female hepatic stellate cell-specific acid ceramidase knockout mice demonstrate no metabolic differences compared to their wildtype counterparts**. (A) Experimental schematic. Female hepatic stellate cell-specific knockout (cACKO) or control (Ctrl) mice were fed either a normal diet (ND) or Fructose Palmitate Cholesterol (FPC) diet for 16 wk. (B) Measured hepatic triglycerides. (C) Liver to body proportional weight in percent. (D-E) Glucose tolerance test was performed at 14 wk and area under the curve (AUC) was determined. (F) Measured serum insulin levels. n = 3-5 mice per group. ***p<0.001.

**Supplemental Figure 2. Ceramide species were analyzed in male hepatic stellate cell-specific acid ceramidase knockout mice or littermate controls in the FPC model of NASH**. Hepatic ceramide subspecies. n = 3-4 mice per group. *p<0.05, **p<0.01, ***p<0.001, ****p<0.0001.

**Supplemental Figure 3. Ceramide species were analyzed in male wild-type mice receiving the FPC diet and B13 treatment**. Hepatic ceramide subspecies. n = 4-5 mice per group. *p<0.05, **p<0.01, ***p<0.001, ****p<0.0001.

