## Supplementary figures and images for "Targeting Acid Ceramidase Ameliorates Fibrosis in Mouse Models of Nonalcoholic Steatohepatitis"

### Supplemental Figure 1

#### Supplemental Figure 1

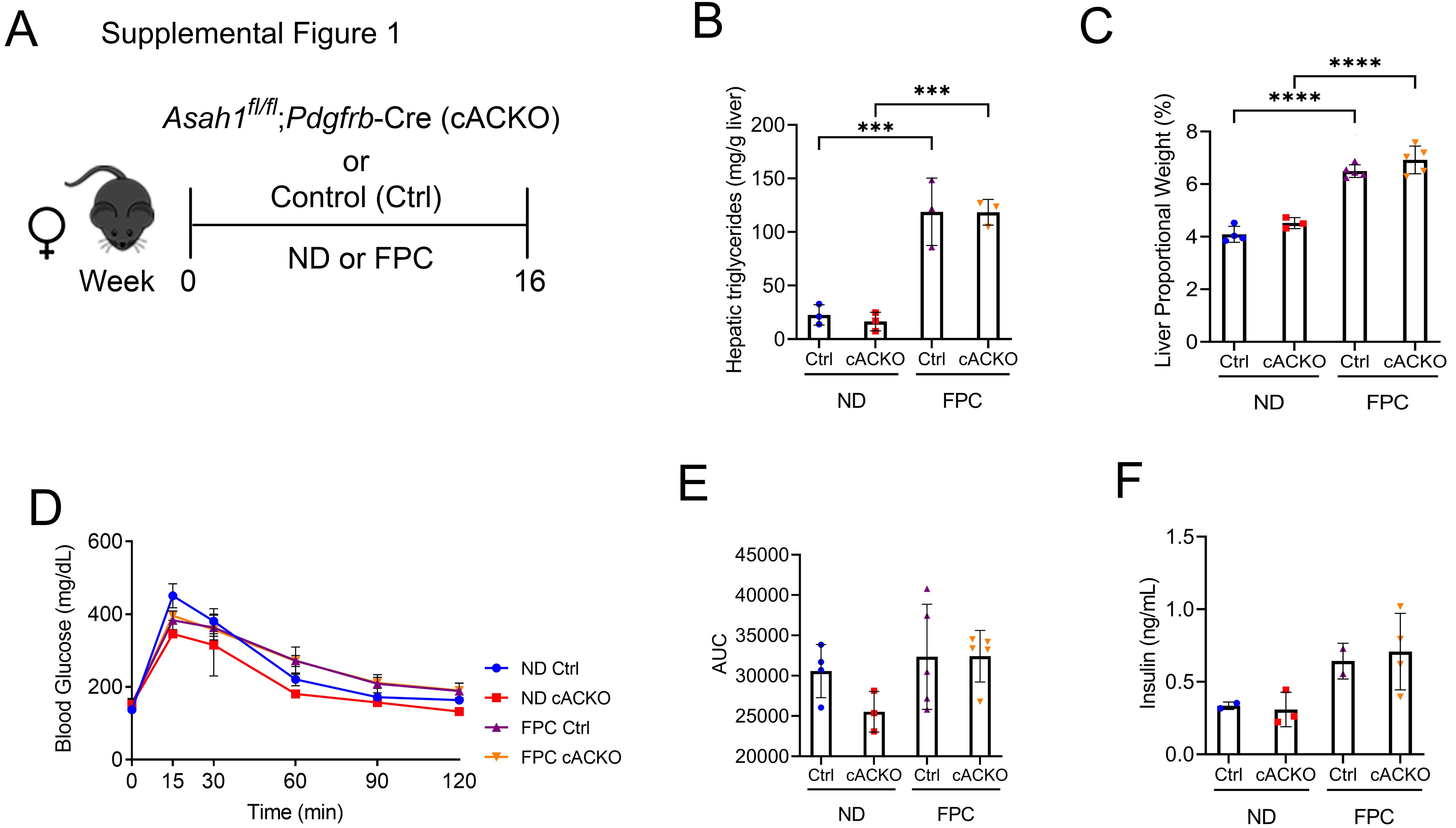

### Supplemental Figure 2

Supplemental Figure 2

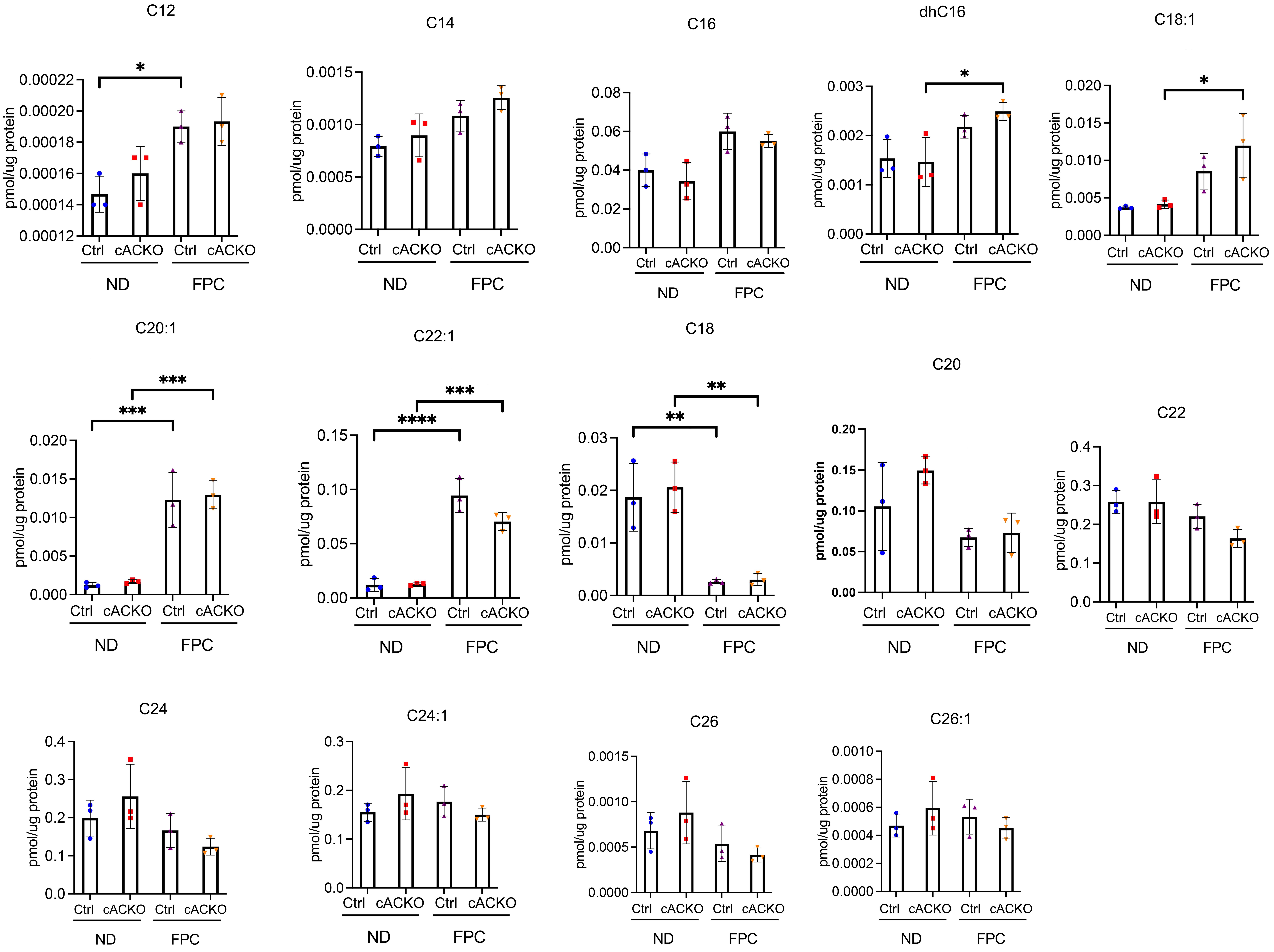

### Supplemental Figure 3

Supplemental Figure 3

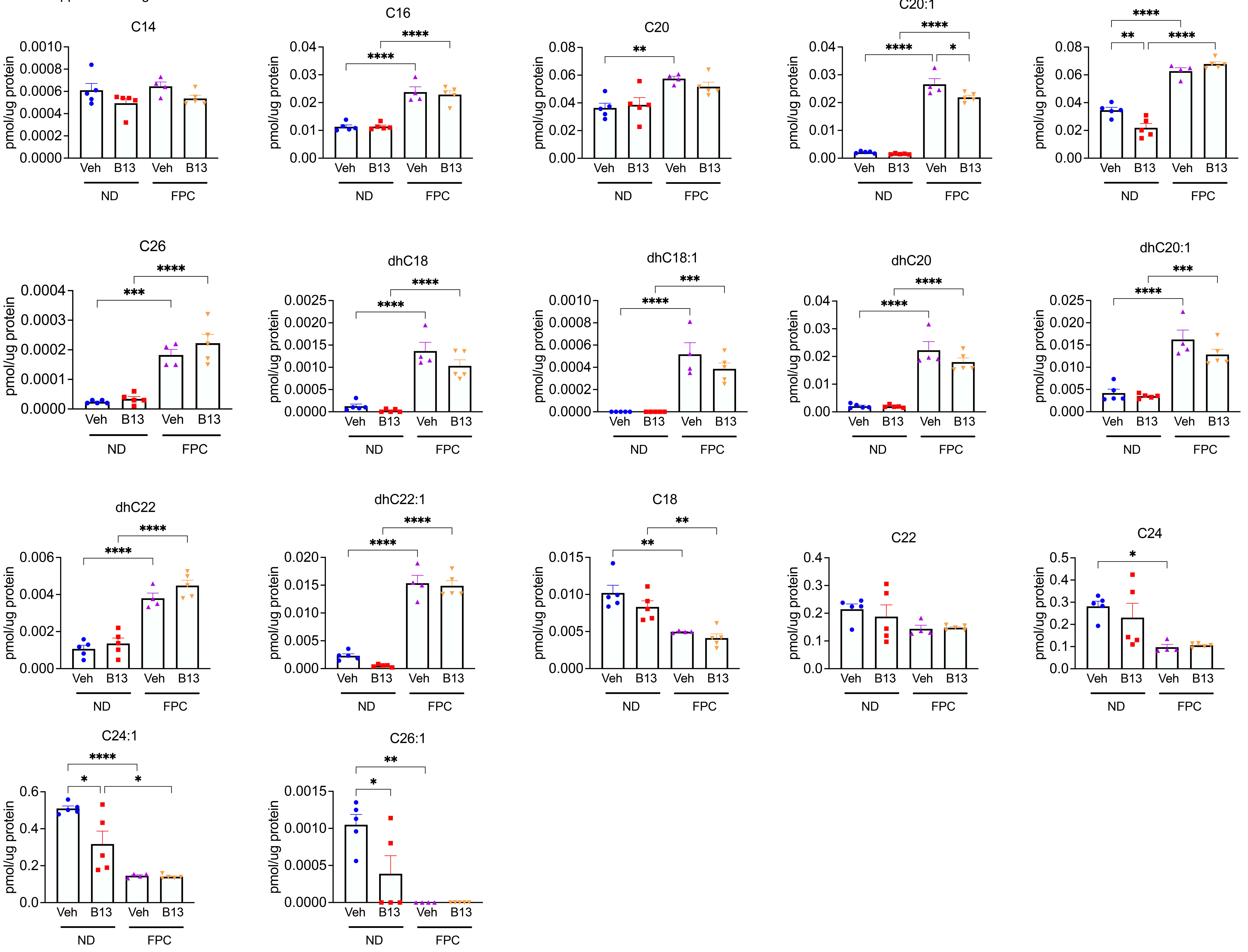
